# Relocating agriculture could drastically reduce humanity’s ecological footprint

**DOI:** 10.1101/488841

**Authors:** Robert M. Beyer, Andrea Manica, Tim T. Rademacher

## Abstract

Agriculture is a major driver of global biodiversity loss, accounts for one quarter of greenhouse gas emissions, and is responsible for 70% of freshwater use. How can land be used for agriculture in a way that minimises environmental impacts while maintaining current production levels? We solved this more than 10 million dimensional optimisation problem and find that moving current croplands and pastures to optimal locations, while allowing then-abandoned areas to regenerate, could simultaneously decrease the current carbon, biodiversity and water footprint of global agriculture by up to 71%, 91% and 100%, respectively. This would offset current net CO_2_ emissions for half a century, massively alleviate pressures on global biodiversity and greatly reduce freshwater shortages. Reductions of up to 59%, 78% and close to 100% are achievable by relocating production within national borders, with the greatest potential for carbon footprint reduction held by the world’s top three CO_2_ emitting countries.

---

The conversion of almost half of the world’s ice-free land area (1) to cropland and pasture has contributed to three of humanity’s most pressing environmental challenges (2, 3): (i) agriculture accounts for a quarter of anthropogenic greenhouse gas emissions (4), largely from the release of carbon stored in vegetation and soils (5, 6); (ii) agriculture is the predominant driver of habitat loss, the greatest threat to global biodiversity (7); and (iii) agriculture is responsible for 70% of global freshwater usage for irrigation, leading to shortages of potable water in many arid areas of the world (8). A rising demand for animal products (9) thwarts hopes that the potential for dietary shifts to decrease the environmental footprints of food production (2, 10, 3) can be fully realised in the near future. Yield increases through more resource-efficient practices, technological advancements and genetically enhanced crop varieties are promising (2, 11, 3), however a growing human population and increasing per-capita consumption (12) threaten to offset the potential of these developments without complementary measures.

Optimising the spatial distribution of production could help to minimise the impact of agriculture (13). Empirical evidence shows that biodiversity and carbon stocks previously lost through land conversion can rapidly reach pre-disturbance levels if these lands are allowed to regenerate, often without active human intervention (Supplementary text). Relocating croplands and pastures that are currently situated in areas with high potential biodiversity and carbon stocks, and subsequently allowing these areas to regenerate, may therefore lead to net carbon and biodiversity benefits. If, in addition, new agricultural areas were established where sufficient rainfall obviates the need for irrigation, the water footprint of global agriculture could be substantially reduced at the same time.

We used global maps of the current distribution of pasture and harvested areas of 43 major crops (Table S1), which between them account for over 95% of global agricultural land (14), to assess the current carbon and biodiversity footprints of agriculture. The carbon impact in a specific area was calculated as the difference between local potential natural carbon stocks in vegetation and soils, and carbon stocks under the type of agricultural land use present there (14). Similarly, the local biodiversity impact of agriculture was estimated as the difference between local biodiversity under natural vegetation, and under cropland or pasture. Biodiversity is measured in terms of range rarity, in which local bird, mammal and amphibian species richness is weighted by the inverse of the species’ ranges (14). Range rarity has been advocated as a particularly meaningful biodiversity metric for conservation planning (15).

By the same methods, we predicted potential carbon and biodiversity impacts in areas that are currently not cultivated but are suitable for agricultural use (14). We used estimates of agro-climatically attainable crop and grass production on potential agricultural areas that assume only rain-fed water supply (14), so as to identify land use configurations that require no irrigation. We considered three different management levels, representing the range from traditional, subsistence-based organic farming systems to advanced, fully mechanised production that uses high-yielding crop varieties and optimum fertiliser and pesticide application (14).

Using these realised and potential yield and impact estimates, we identified the global distribution of agricultural areas that provides the same total production of the 43 crops and grass as the current one, while minimising the total environmental footprint. On a 30 arc-minute (0.5°) grid, this requires solving a more than 10^6^-dimensional linear optimisation problem (14). We estimated that for the optimal configuration of agricultural areas and advanced management farming, current carbon and biodiversity impacts of global agriculture could be simultaneously reduced by up to 71% and 91%, respectively (Fig. 1A). This would offset the current annual increase of atmospheric CO_2_ of 4.7 Pg C y^−1^ (16) for 49 years, while drastically alleviating pressures on terrestrial biodiversity. As per the data used, no irrigation is required to supplement rainfall water supply. The total worldwide area used for agriculture in this scenario is less than half of its current extent. The trade-off between reducing carbon and biodiversity impacts is minimal; optimising land use for each impact measure independently yields only marginally higher reduction potentials of 74% and 98%, respectively (Fig. 1A). Under traditional farming, simultaneous carbon and biodiversity impact reductions of up to 43% and 84%, respectively, are feasible (Fig. 1C). Whilst this confirms that increasing crop yields is important for reducing the environmental footprint of agriculture (11, 2, 12, 10, 3), it demonstrates that a substantial impact reduction could be achieved by land reallocation alone.

**Figure 1:**
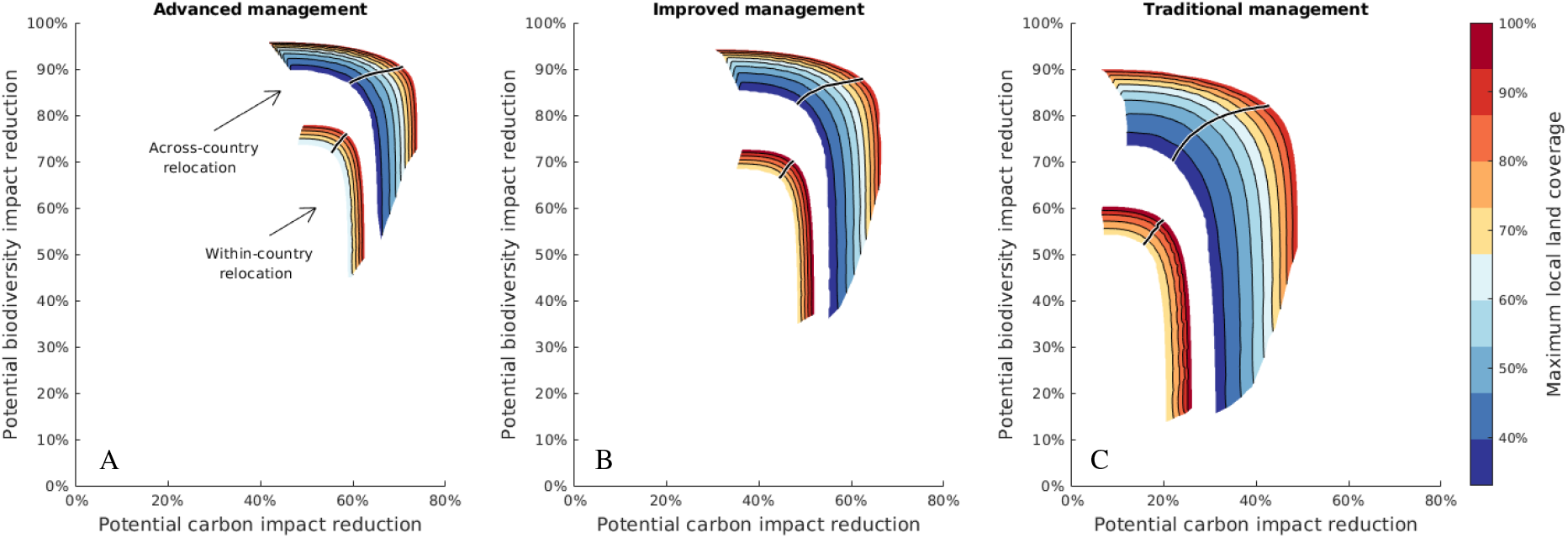
Possible reduction of current carbon and biodiversity footprints of global agriculture by optimally relocating agricultural production. Each contour level represents the frontier of simultaneously achievable carbon and biodiversity impact reduction for a given proportion of local land area that is available for agricultural use (see text). Black lines correspond to a subjectively defined optimal trade-off between carbon and biodiversity impacts.

Thus far, we have assumed that the entire area of each grid cell is available for agricultural use. How does the potential for reducing impact change if only a proportion of each grid cell can be cultivated, while the remainder is retained as natural ecosystem or used for other purposes? In this scenario, total impacts are necessarily higher, because less optimal areas, in which environmental impacts are higher in relation to yields, also need to be cultivated to meet a given production level. This is disproportionally the case for low intensity farming, which inherently requires more land. We found that when only half of the local land area can be used for agriculture, carbon and biodiversity impacts could be simultaneously reduced by 63% and 90%, respectively, under advanced management (Fig. 1A), but only by 30% and 80%, respectively, for traditional farming (Fig. 1C). Allocating as much land as possible in optimal areas therefore becomes more important the less advanced the farming system is.

Moving agricultural production, and thus labour and capital, across national borders poses political and socio-economic challenges that will be difficult to resolve in the near future. We therefore repeated our analyses, allowing croplands and pastures to be relocated only within countries, while requiring current national production levels to remain unchanged (14). We estimated that if each country independently optimised its distribution of agricultural areas, the current global carbon and biodiversity impacts of agriculture could be simultaneously reduced by up to 59% and 78%, respectively (Fig. 1A). In this scenario, the vast majority of production can be relocated so that rainfall provides sufficient water supply; however, some countries produce crops for which national natural agro-climatic conditions are not suitable, and thus some irrigation continues to be needed (14). Fig. 3 lists the ten countries with the highest absolute carbon and biodiversity reduction potentials, showing that the world’s three largest CO_2_ emitters – China, India and the United States (17) – are also the countries that can reduce their agricultural carbon footprint the most.

Agricultural areas optimally sited to minimise environmental impacts coincide only to a limited extent with their current distribution (Fig. 2). The world’s most produced crop, maize, for example, is currently planted predominantly in the United States and China, but would ideally be grown in parts of Sub-Saharan Africa (Fig. S1). In the scenario of optimal within-country land reallocation, the optimal land use coincides with the current one on 30% of optimal areas, while 42% of optimal areas are located in regions already under some type of agricultural use, and 45% are located in either currently active or abandoned agricultural areas (Fig. S2). This overlap is significantly lower in the scenario of across-country relocation (Fig. S2). Whilst the expansion of agriculture into degraded areas has been advocated as a way to minimise future biodiversity and carbon losses (2, 11), our results suggest that potential biodiversity and carbon stocks on currently cultivated and abandoned agricultural areas are often so high that, in principle, their restoration would be preferable to the protection of natural habitat in the identified optimal growing areas.

**Figure 2:**
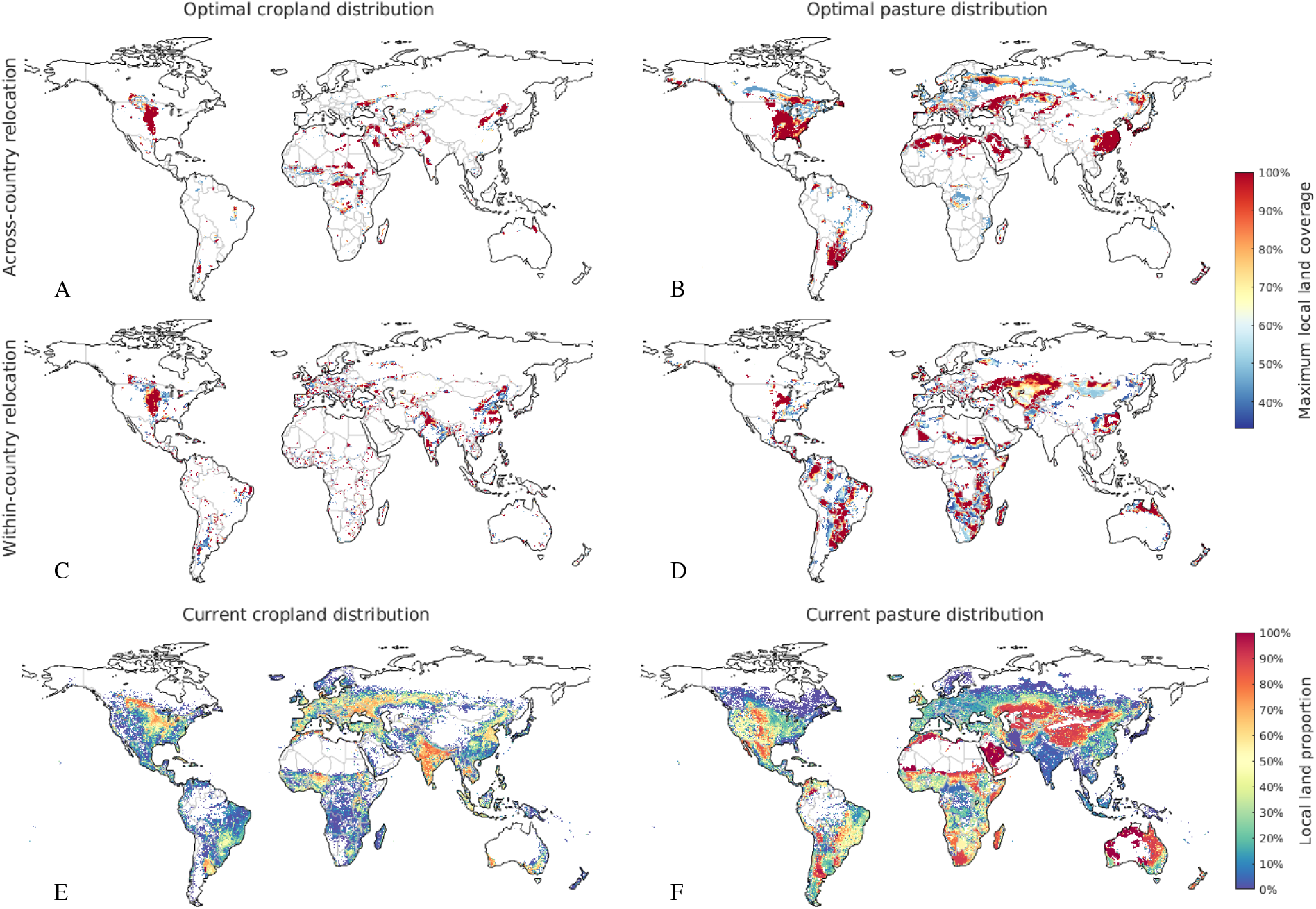
Optimal and current distribution of agricultural areas. **(A)–(D)** Optimal distribution of croplands and pastures for across- and within-country relocation of current agricultural areas. Red areas represent the land that is required to meet current production levels if grid cells are entirely available for agricultural use, while both red and yellow areas are required when only 50% of grid cells are allowed to be cultivated, etc. (see text). Maps show optimal configurations for advanced management farming and the optimal impact trade-off shown in Fig. 1A. **(E)–(F)** Current distribution of croplands and pastures. Current and optimal distributions for a specific crop, maize, are shown in Figure S2C–E.

**Figure 3:**
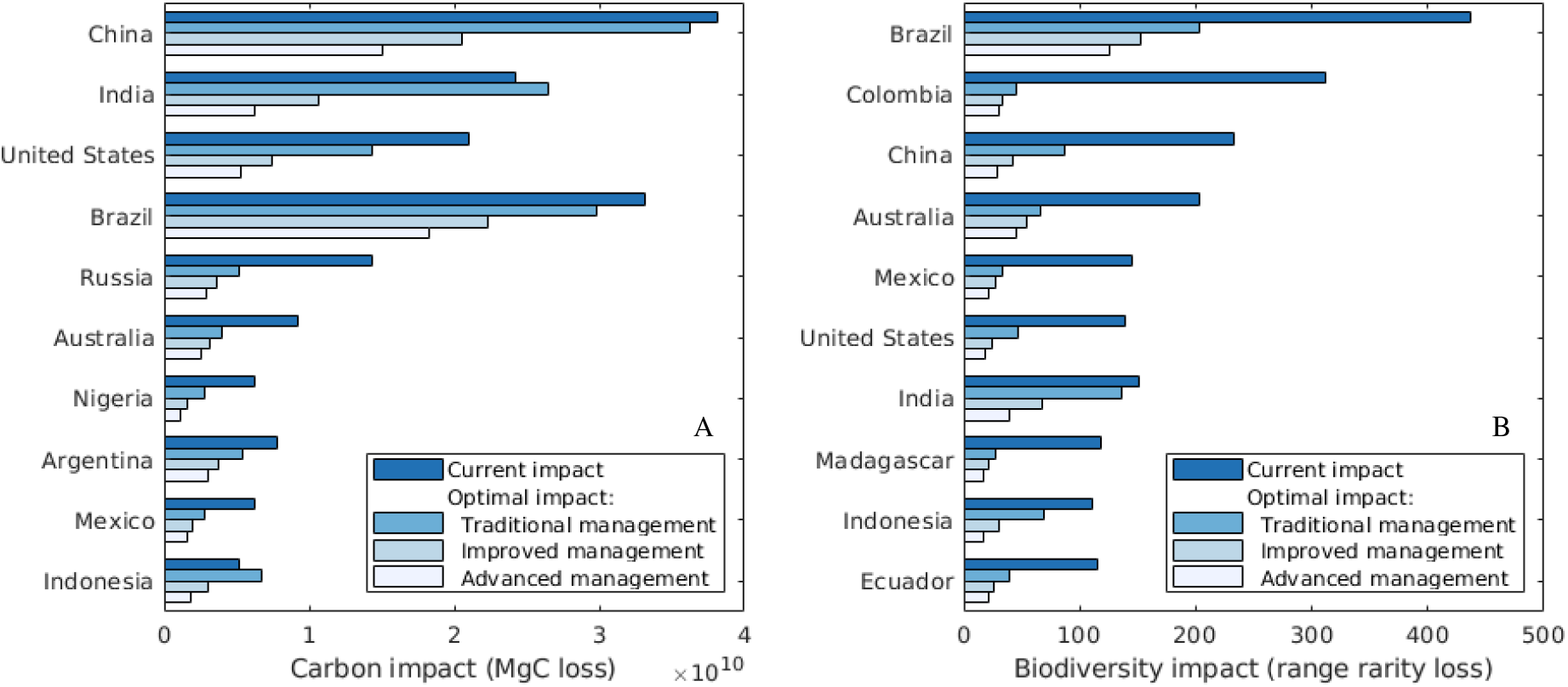
Current national **(A)** carbon and **(B)** biodiversity impacts of agriculture, and potentials for impact reduction by means of within-border relocation of agricultural areas for 10 countries, ordered by absolute impact reduction potential. Results correspond to the optimal impact trade-offs shown in Fig. 1A–C.

For computational reasons, here, we did not explore the possibility of crop rotations and other diversification types, which can have benefits, e.g. for pest and disease suppression (18). Our analyses also do not account for possible changes in yields as the result of climate change (19). Both aspects may affect the precise location of optimal areas, and decrease or increase the reduction potentials identified here, however, we do not expect them to qualitatively change our overall conclusions.

Whilst our estimates of achievable impact reductions assume a fully optimised distribution of agricultural areas, we stress that even relocating only a small share of production would already generate a substantial portion of these benefits. 50% of the current total carbon impacts of individual crops are caused by areas that account for only 26±5% of the total production, while a mere 8±4% of production are responsible for 50% of biodiversity impacts (Fig. S3). Prioritising the relocation of these areas, where the ratio of environmental impact to yield is largest, would have disproportionately high carbon and biodiversity benefits, and represents the most effective strategy for countries to progressively reduce impacts.

How can the relocation of agricultural areas, whether at a small or large scale, be implemented in practice? There are a number of national and supranational set-aside schemes, aimed at retiring agricultural land for environmental benefits, that offer useful templates for how direct payments for ecosystem services can reduce impacts in socio-economically sustainable ways (20, 21). As a case example, China’s Grain for Green programme, the world’s largest national set-aside scheme, achieved the reforestation of 15 million hectares of farmland between 1999 and 2010, with generally positive economic outcomes for the 124 million people involved (22). While primarily aimed at reducing soil erosion, the programme also generated substantial carbon and biodiversity benefits (22, 23). Notably, the scheme led to an effective relocation of agricultural land from Southern to Northern China (24), facilitated by higher financial incentives for retiring land in the South. Setting up incentives in relation to local potential biodiversity and carbon stocks will indeed be crucial for achieving benefits most effectively. International climate funds can support countries that lack the financial means for direct payments to farmers in implementing durable set-aside schemes. It is important to note that in many parts of the world, agricultural subsidies prevent land abandonment and migration to urban centres that would naturally occur otherwise (25); thus, reducing subsidies in areas with high potential biodiversity and carbon stocks represents a particular cost-efficient way to generate benefits (26). A range of financial, infrastructural and policy measures have also proven effective at steering the establishment of new agricultural land towards desired agro-ecologically optimal areas (27). Spatial reallocation of agricultural production has tremendous potential to reduce its environmental footprint, but the implementation of such changes requires careful management of the process. Relocating cultivated areas can only lead to a reduction of impact if abandoned areas with high potential biodiversity and carbon stocks are effectively protected and their regeneration is ensured. This requires strong institutional, legal, and policy frameworks, and financial incentives for landowners (28, 29, 30). Their implementation at the national and international level will be crucial for realising the environmental potential of moving agricultural areas, providing gains that are badly needed if we are to reverse the ongoing degradation of global climate, biodiversity and water under an ever increasing demand for food.

## Acknowledgments

We thank Günther Fischer, Paul Donald, Philip Martin and Fangyuan Hua for their advice and comments during the preparation of this manuscript. This work benefited from conversations with Fiona Sanderson, Catherine Tayleur, Paul Donald, Sharon Brooks, David Coomes, América P. Durán and Phil Martin during a separate research project that was supported by a grant from the Cambridge Conservation Initiative Collaborative Fund (CCI-06-16-008).

## Author contributions

R.M.B. designed the project, did the analysis and wrote the manuscript. All authors interpreted the results and commented on the manuscript.

## Materials and Methods

In the following, we define the mathematical optimisation problem whose solutions represent minimum impact configurations of agricultural land, and specify the datasets that were used to solve it. We use the following notation:

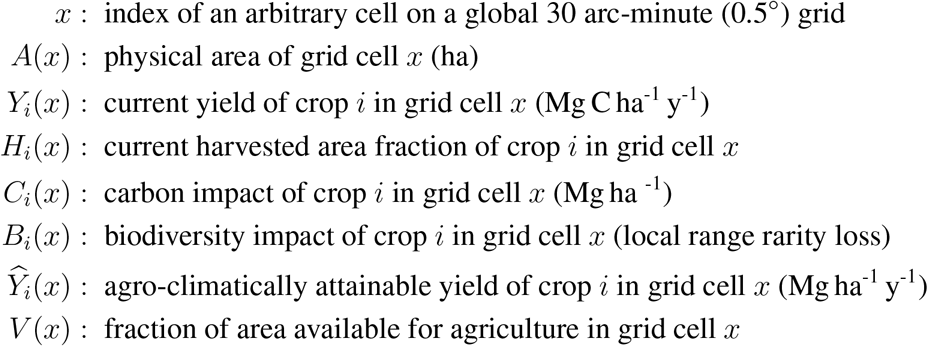

On pastures, yield is assessed in terms of the annual production of forage per hectare. The current total annual production of crop *i* is given by

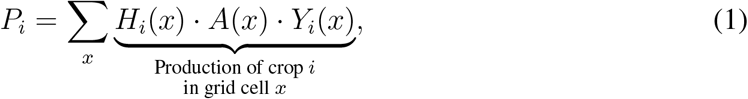

and the current global carbon and biodiversity impacts of agriculture are given by

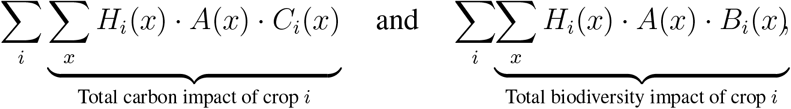

respectively.

For each crop *i* and each grid cell *x*, we determined the harvested area fraction *Ĥ*_*i*_(*x*) such that the total production of each crop *i* equals the current production *P*_*i*_, while the environmental impact is minimised. Any solution must satisfy the equality constraints

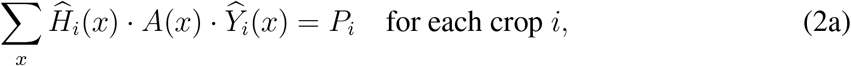

requiring the total production on new agricultural areas to be equal to the current one, and the inequality constraints

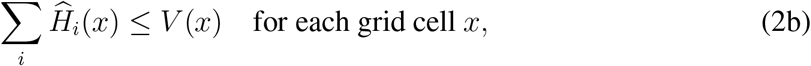

which ensure that the local sum of agricultural lands is not larger than the locally available area.

Subject to these constraints, we can identify the configuration that minimises the total carbon or biodiversity impact by minimising the objective function

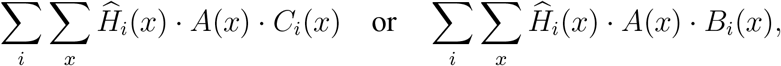

respectively. More generally, we consider the linearly weighted objective function

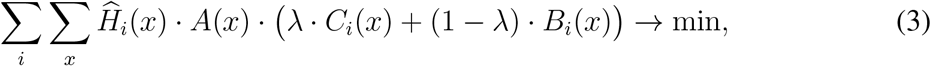

where *λ* ranges between 0 and 1, thus allowing us to minimise both impacts simultaneously and examine potential trade-offs.

The above framework is identical when examining the potential for impact reduction by means of relocating croplands within national borders rather than globally. In this case, the sum over *x* in the calculation of national production (Eq. (1)), in the optimisation constraints (Eqs. (2a)–(2b)) and in the objective function (Eq. (3)) is taken over grid cells that correspond to specific countries rather than the whole world, and the optimisation problem is solved independently for each country. Some countries produce small quantities of crops that, according to the data used here, would not grow anywhere within their borders under natural climatic conditions, i.e. these crops likely require irrigation or greenhouses cultivation. Our analysis shows that these crops account for a fraction of 0.12% of current global agricultural areas that can not be relocated within national borders to areas where rain-fed cultivation is possible. These crops were excluded from Eq. (3) for the respective countries; we added the environmental impacts associated with the current growing areas of these crops to the minimum national impacts found by Eq. (3).

Although all data required to compute the relevant variables, *A*(*x*), *Y*_*i*_(*x*), *H*_*i*_(*x*), *C*_*i*_(*x*), *B*_*i*_(*x*), *Ŷ*_*i*_(*x*) and *V* (*x*) (see below), are available at a 5 arc-minute (0.083°) grid resolution, for computational reasons, we upscaled the final data to a 30 arc-minute (0.5°) grid. For pasture and 43 crops, this implies a more than 10 million dimensional linear optimisation problem. We solved Eq. (3) using the dual-simplex algorithm in the function *linprog* of the Matlab R2018a Optimization Toolbox (31).

### Current and potential agricultural areas and yields

**H**_**i**_(**x**), **Y**_**i**_(**x**), **Ŷ**_**i**_(**x**) We used global maps of harvested areas, *H*_*i*_(*x*), and fresh weight yields, *Y*_*i*_(*x*), of 43 crops (32), and a global map for pasture (33) (Table S1). These areas cover 95.2% of the combined area of pasture and harvested areas of 175 crops (32), for which data is available. We used global maps of potential growing areas and agro-climatically attainable dry weight yields, *Ŷ*_*i*_(*x*), for baseline climate, rain-fed water supply and three different management levels for the same 43 crops and pasture grass (34). Management levels represent the range from traditional, labour-intensive farming systems without synthetic chemicals, to advanced, market-oriented production that is fully mechanised, uses high-yielding crop varieties, and optimum applications of nutrients and pest, disease and weed control (34). Potential yields were converted from dry weight to fresh weight using crop-specific conversion factors (32). We are not aware of a global dataset of forage production on current pastures, and therefore used potential pasture grass yields for rainfed water supply and intermediate input management as an estimate on these areas.

### Carbon impact

**C**_**i**_(**x**) Following ref. (6), the local carbon impact of agriculture, *C*_*i*_(*x*), was estimated as the difference between potential natural vegetation and soil carbon stocks, and carbon stocks under agricultural land cover.

The change of carbon stocks in vegetation resulting from land conversion is given by the difference of carbon stored in potential natural vegetation (6) and carbon stored in grass or crops, which was calculated as in ref. (6), based on the data compiled by ref. (32).

Due to the technical difficulties of acquiring empirical data across large spatial scales, spatially-explicit global estimates of soil organic carbon (SOC) dynamics under varying land use types are currently not available. We therefore chose a simple approach, consistent with average estimates across large spatial scales, rather than a complex spatially-explicit model for which, given the limited empirical data, robust predictions on and beyond currently cultivated areas would not be possible. Following ref. (6), and supported by empirical meta-analyses (35, 36, 37, 38, 39), we assumed a 25% reduction of potential natural SOC (see below) from the conversion to cropland. Meta-analyses of the change of SOC stocks when natural habitat is converted to pasture suggest, on average, no significant change (37), a slight increase (36, 39) or slight decrease (38). Here, we assumed no change in carbon stocks when natural habitat is converted to pasture. Absolute local SOC loss from the conversion of potential natural vegetation to cropland or pasture was estimated by applying the appropriate loss percentages to a global map of pre-agricultural SOC stocks (5). The total local carbon impact of agriculture (Mg C ha^−1^) is thus given by

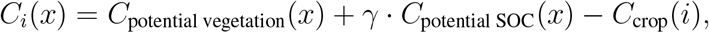

where *C*_potential vegetation_(*x*), *C*_potential SOC_(*x*) and *C*_crop_(*i*) denote the carbon stocks (Mg C ha^−1^) of potential natural vegetation, potential natural SOC stocks, and carbon stocks of crop *i*, respectively, in the grid cell *x*, and where *γ* is equal to 0.25 or 0 if land is converted to cropland or pasture, respectively.

We did not consider greenhouse gas emissions from sources other other than from land use change. This includes nitrous emissions from fertilised soils and methane emissions from livestock and rice paddies (40). In contrast to the one-off land use change emissions, these are ongoing emissions that are tied to production and incur continually. We do not consider available data sufficient to allow a robust extrapolation of these emission types to currently uncultivated land. We argue, though, that the magnitude of these emissions in a scenario of land reallocation in which total production is constant, is likely similar to that associated with the current distribution of agricultural areas. We also did not consider emissions associated with transport; however, these have been argued to be small compared to other food chain emissions (41) and poorly correlated with the actual distance travelled by agricultural products (42).

### Biodiversity impact

**B**_**i**_(**x**) We assessed the local biodiversity impact of agriculture in terms of range rarity loss. Range rarity has been advocated as a metric for biodiversity that is more relevant to conservation planning than alternative measures, such as species richness (43, 44, 45, 15, 46). *B*_*i*_(*x*) is calculated as the difference between range rarity under natural vegetation and under agricultural land cover as follows: Using a similar approach to that of ref. (47), we considered a bird, mammal or amphibian species to be potentially present in a cell of a 5 arc-minute grid if the species’ spatial extent of occurrence (48, 49) overlays the grid cell, and if its habitat preferences (48, 49) include the local potential natural vegetation type (50). Each species’ potential natural range (ha) is then given by the total area of all grid cells identified as containing the species. Next, potential natural range rarity of each grid cell was obtained as the sum of the inverse ranges of all species present in the grid cell under potential natural vegetation. Finally, global maps of range rarity loss resulting from the conversion of natural vegetation to cropland or pasture were derived by subtracting, in each grid cell, the sum of the inverse ranges of potentially present species whose habitat preferences also include cropland or pasture, respectively, from the potential natural range rarity. As with *C*_*i*_(*x*) (see above), this approach allowed us to estimate biodiversity impact for both currently cultivated and uncultivated areas.

### Land available for agriculture

**V**(**x**) We assumed that the maximum area available for agriculture in a grid cell is given by the proportion not occupied by any crop other than the 43 considered here (32), or by water bodies, infrastructure or settlements (34). Areas where soil and terrain-slope conditions are not suitable for agriculture are already excluded in the potential yield data (34).

As specified in the main text, we also examined the scenario in which only a certain fraction of this maximum available area is available as potential agricultural land.

**Figure S1:**
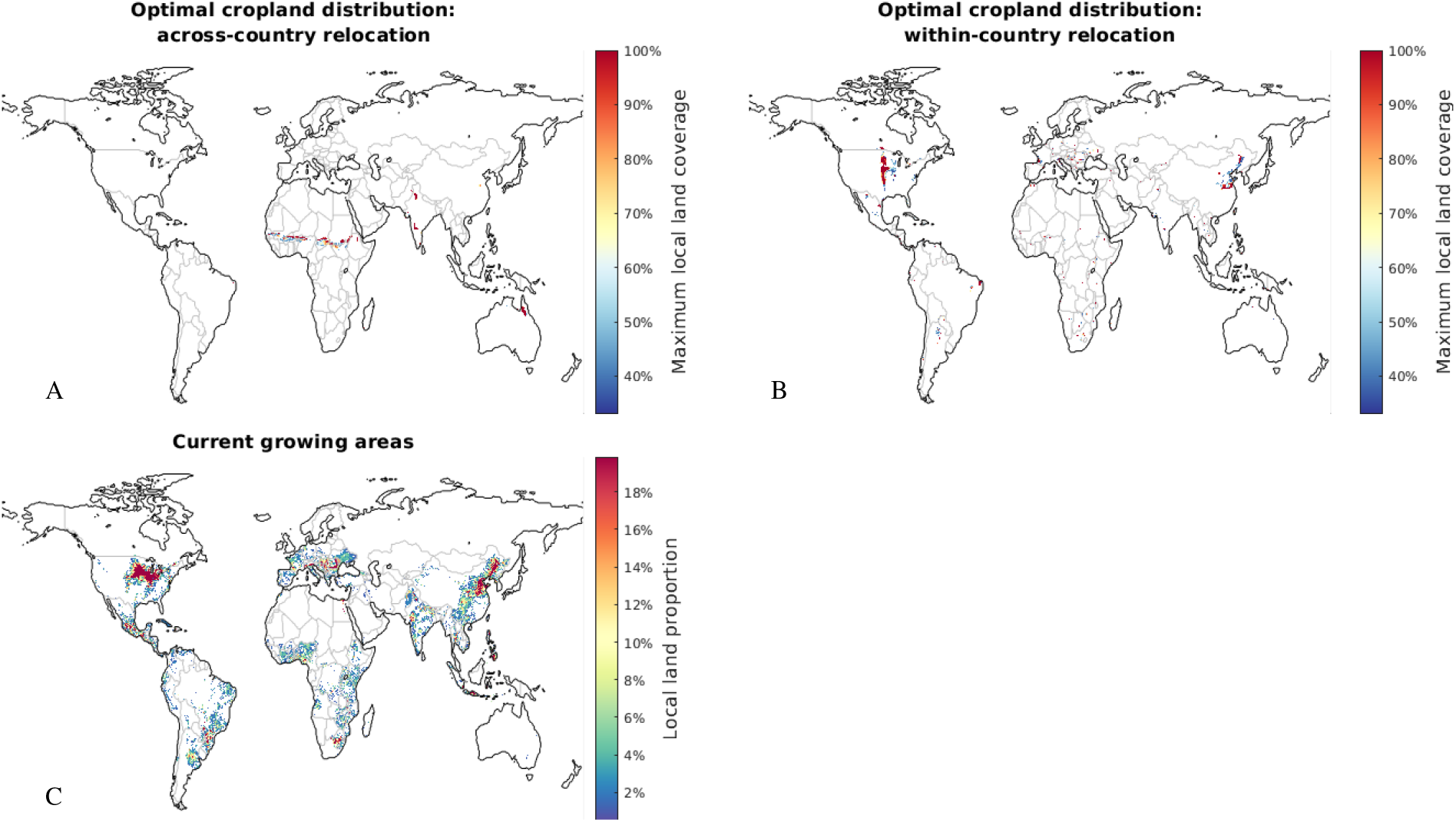
**(A)–(C)** Optimal and current growing locations of maize in the scenarios shown in Fig. 1A,C,E, respectively.

**Figure S2:**
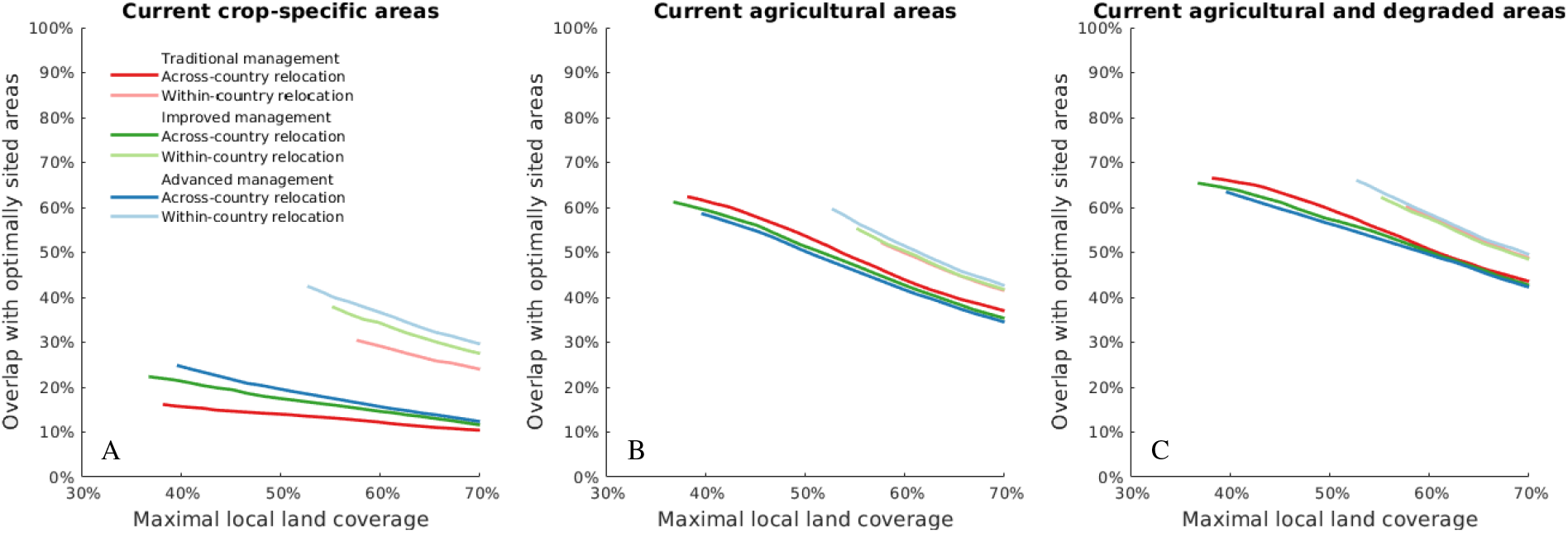
Overlap between optimal agricultural areas (under the six specified scenarios) and **(A)** current agricultural land where the specific current and optimal crop types coincide (i.e. where current crops are already optimally sited), **(B)** current agricultural land (irrespective of the specific local land use), and **(C)** the combined area of current agricultural land and abandoned, wasted or idle agricultural land (using the global dataset of ref. (51)). All values are relative to the total area required in the appropriate optimal scenario. Plots correspond to the optimal impact trade-off shown in Fig. 1A–C.

**Figure S3:**
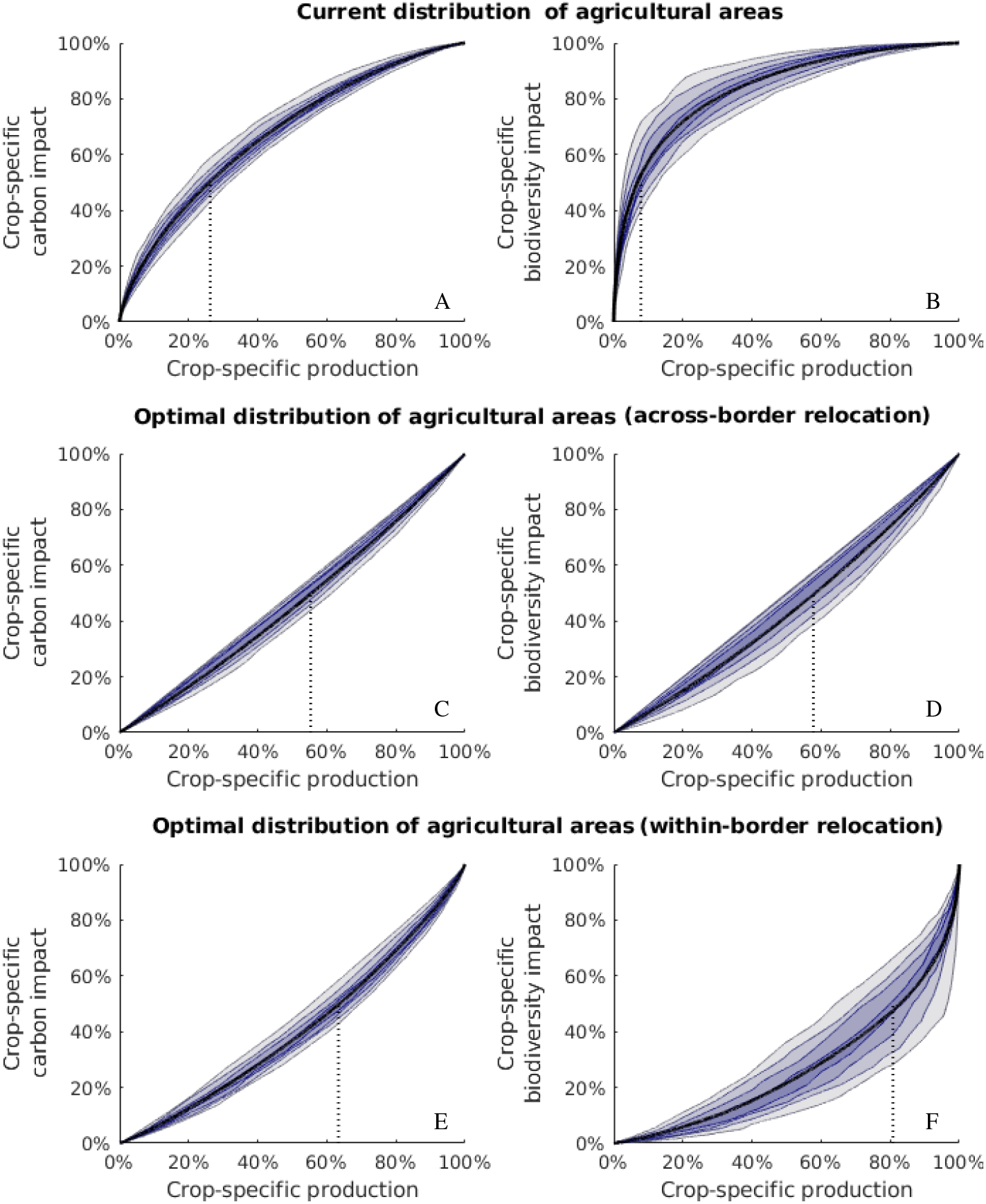
Cumulative production and environmental impact on current and optimal agricultural areas. **(A)–(B)** For each crop and grass, we sorted the relevant growing locations according to the local ratio of environmental impact to yield, from high to low. We then generated cumulative production-impact curves by traversing areas in that order. For comparability, the resulting 44 curves were converted to a relative scale, and are summarised by the 10–90th percentiles shown. Black lines represent means across crops. Dotted lines illustrate the level of production on the least agro-environmentally efficient growing areas that corresponds to 50% of the total crop-specific impact. The more concave the production-impact curve is, the larger are the relative environmental benefits of relocating even small portions of land. **(C)–(F)** Equivalent of (A)–(B) for optimally sited agricultural areas, but with growing locations ordered from low to high impact-to-yield ratio (i.e. the order in which new areas would ideally be established). Dotted lines thus show the maximum achievable level of production that causes 50% of the environmental impact. Data correspond to advanced management farming and the optimal impact trade-off shown in Fig. 1A.

**Table S1:** List of crops considered in this study, their current global carbon and biodiversity impacts, and the potentials for reducing impacts under optimal within- and across-border relocation of areas. Reduction potentials are shown for advanced management farming and the optimal impact trade-off shown in Fig. 1A.

|  | Current carbon impact of agriculture (Mg C) | Current biodiversity impact of agriculture (cumulative range rarity) | Potential national carbon impact reduction | Potential international carbon impact reduction | Potential national biodiversity impact reduction | Potential international biodiversity impact reduction |
| --- | --- | --- | --- | --- | --- | --- |
| Alfalfa | 21.9 | 8.3 | 77% | 87% | 83% | 83% |
| Banana | 8.3 | 23 | 64% | 74% | 82% | 97% |
| Barley | 69.5 | 18.5 | 78% | 90% | 87% | 79% |
| Buckwheat | 3.4 | 0.9 | 76% | 80% | 84% | 83% |
| Cabbage | 4.5 | 3.1 | 73% | 90% | 80% | 93% |
| Cassava | 30.7 | 46.1 | 77% | 90% | 90% | 95% |
| Carrot | 1.5 | 1.1 | 69% | 87% | 87% | 88% |
| Chickpea | 14.2 | 8.6 | 66% | 92% | 86% | 96% |
| Cocoa | 14.3 | 27.7 | 87% | 92% | 95% | 99% |
| Coconut | 26.2 | 86 | 68% | 81% | 87% | 97% |
| Cocoyam | 2.8 | 4.2 | 82% | 88% | 94% | 97% |
| Coffee | 22.2 | 72.4 | 85% | 89% | 96% | 99% |
| Cotton | 33.7 | 18.8 | 58% | 80% | 80% | 79% |
| Cowpea | 9.9 | 6.6 | 92% | 94% | 94% | 94% |
| Flax | 0.8 | 0.1 | 47% | 70% | 58% | 39% |
| Green bean | 1.6 | 2 | 89% | 93% | 97% | 99% |
| Groundnut | 30.7 | 21.9 | 80% | 89% | 89% | 92% |
| Maize | 202.4 | 225.4 | 76% | 84% | 92% | 92% |
| Millet | 30.2 | 18.6 | 91% | 95% | 92% | 96% |
| Oat | 17.9 | 4.1 | 68% | 84% | 79% | 74% |
| Oil palm | 24.1 | 42.6 | 58% | 71% | 81% | 93% |
| Olive | 9.9 | 3.1 | 87% | 96% | 95% | 95% |
| Onion | 4.3 | 3.6 | 78% | 90% | 91% | 91% |
| Orange | 6 | 9.4 | 74% | 86% | 89% | 96% |
| Pasture grass | 97.8 | 24.4 | 83% | 95% | 89% | 86% |
| Pasture legume | 6.3 | 1.9 | 49% | 80% | 85% | 64% |
| Pea | 8 | 3.2 | 58% | 72% | 73% | 86% |
| Pigeon pea | 6.9 | 3.7 | 91% | 94% | 87% | 89% |
| Potato | 31.2 | 17.6 | 77% | 90% | 84% | 94% |
| Rape | 34.9 | 11.1 | 73% | 89% | 79% | 87% |
| Rice | 308.6 | 305.1 | 62% | 75% | 84% | 91% |
| Rye | 15.2 | 1.2 | 75% | 85% | 77% | 57% |
| Sorghum | 39.6 | 33.2 | 92% | 92% | 94% | 96% |
| Soybean | 105 | 56 | 73% | 75% | 78% | 85% |
| Sugarbeet | 9.2 | 1.2 | 52% | 71% | 67% | 51% |
| Sugarcane | 36.1 | 67.2 | 20% | 55% | 62% | 87% |
| Sunflower | 23.2 | 8.2 | 76% | 83% | 83% | 91% |
| Sweet potato | 15.6 | 15.8 | 66% | 88% | 93% | 92% |
| Tea | 4.2 | 9.9 | 75% | 85% | 75% | 97% |
| Tobacco | 7.7 | 8.3 | 68% | 86% | 86% | 96% |
| Tomato | 5.1 | 4.1 | 61% | 83% | 81% | 76% |
| Wheat | 244.4 | 79.8 | 74% | 86% | 68% | 88% |
| Yam | 7.1 | 5.8 | 81% | 85% | 85% | 92% |
| Pasture | 1807.3 | 2808.9 | 48% | 59% | 75% | 90% |

## Supplementary text

### Carbon and biodiversity recovery on abandoned agricultural land

We briefly review empirical results on the recovery of biodiversity and carbon stocks on regenerating agriculturally degraded land. In both cases, recovery follows an asymptotic concave trajectory; the time required to reach pre-disturbance levels can therefore be difficult to pinpoint, as both slightly shorter or longer times correspond to similar recovery levels in the flat saturation stage of the recovery function. Above- and below-ground carbon stocks have been found to asymptotically reach pre-disturbance values within 50–100 years after land abandonment (52, 53, 54, 55, 56, 57, 58). Slower biomass accumulation coupled with lower potential carbon stocks in temperate forests leads to overall similar recovery times compared to tropical forests (59). In grasslands, carbon stocks recover within a few decades following land abandonment (60, 54). Faunal species richness on regenerating degraded land reaches pre-disturbance levels on timescales of decades to a century (61, 62, 56, 63, 53, 64). Initial colonists may represent different species to those present before degradation occurred (65, 63, 66), but the proportion of old-growth species increases as secondary ecosystems age, thereby gradually replacing non-native species (67, 65, 68, 63, 56, 69, 70). Biodiversity tends to regenerate faster in temperate than in tropical regions, and faster in grassland and shrubland biomes than in forests (62, 71, 53, 64).

Whilst assisted regeneration and active restoration can accelerate carbon and biodiversity recovery (28, 67, 29, 62, 72, 73, 74, 63, 75, 66, 53), passive regeneration is often the most effective strategy (76).

## References

[1] K. Klein Goldewijk, A. Beusen, J. Doelman, E. Stehfest, Earth System Science Data 9, 927 (2017).

[2] J. A. Foley, et al., Nature 478, 337 (2011).

[3] M. Springmann, et al., Nature 562, 519 (2018).

[4] S. J. Vermeulen, B. M. Campbell, J. S. Ingram, Annual Review of Environment and Resources 37, 195 (2012).

[5] J. Sanderman, T. Hengl, G. J. Fiske, Proceedings of the National Academy of Sciences 114, 9575 (2017).

[6] P. C. West, et al., Proceedings of the National Academy of Sciences 107, 19645 (2010).

[7] O. E. Sala, et al., Science 287, 1770 (2000).

[8] S. L. Postel, G. C. Daily, P. R. Ehrlich, Science 271, 785 (1996).

[9] P. K. Thornton, Philosophical Transactions of the Royal Society of London B: Biological Sciences 365, 2853 (2010).

[10] D. Tilman, et al., Nature 546, 73 (2017).

[11] J. Clay, Nature 475 (2011).

[12] D. Tilman, C. Balzer, J. Hill, B. L. Befort, Proceedings of the National Academy of Sciences 108, 20260 (2011).

[13] D. K. Frankel, M. C. Rulli, A. Seveso, P. D’Odorico, Nature Geoscience 10 (2017).

[14] Materials and Methods are available as Supplementary Materials.

[15] G. R. Guerin, A. J. Lowe, Biodiversity and Conservation 24, 2877 (2015).

[16] C. Le Quéré, et al. Earth System Science Data Discussions 2018, 1 (2018).

[17] J. G. Olivier, G. Janssens-Maenhout, M. Muntean, J. A. Peters (2016).

[18] B. B. Lin, BioScience 61, 183 (2011).

[19] C. Rosenzweig, et al., Proceedings of the National Academy of Sciences 111, 3268 (2014).

[20] OECD, Environmental Indicators for Agriculture (2001).

[21] J. Van Buskirk, Y. Willi. Conservation Biology 18, 987 (2004).

[22] C. O. Delang, Z. Yuan, China’s grain for green program (Springer, 2016).

[23] F. Hua, et al., Nature Communications 7, 12717 (2016).

[24] J. Liu, et al., Journal of Geographical Sciences 24, 195 (2014).

[25] S. Li, X. Li, Journal of Geographical Sciences 27, 1123 (2017).

[26] N. Myers, Perverse subsidies: how tax dollars can undercut the environment and the economy (Island Press, 2001).

[27] B. Phalan, et al., Science 351, 450 (2016).

[28] D. Lamb, P. D. Erskine, J. A. Parrotta, Science 310, 1628 (2005).

[29] R. L. Chazdon, Science 320, 1458 (2008).

[30] J. M. Bullock, J. Aronson, A. C. Newton, R. F. Pywell, J. M. Rey-Benayas, Trends in Ecology & Evolution 26, 541 (2011).

[31] MATLAB and Optimization Toolbox Release 2018a.

[32] C. Monfreda, N. Ramankutty, J. A. Foley, Global Biogeochemical Cycles 22 (2008).

[33] N. Ramankutty, A. T. Evan, C. Monfreda, J. A. Foley, Global Biogeochemical Cycles 22 (2008).

[34] G. Fischer, et al., Global Agro-ecological Zones (GAEZ v3. 0)-Model Documentation, Tech. rep., IIASA, Laxenburg, Austria and FAO, Rome, Italy (2012).

[35] R. A. Houghton, Tellus B 51, 298 (1999).

[36] L. B. Guo, R. Gifford, Global Change Biology 8, 345 (2002).

[37] D. Murty, M. U. Kirschbaum, R. E. Mcmurtrie, H. Mcgilvray, Global Change Biology 8, 105 (2002).

[38] A. Don, J. Schumacher, A. Freibauer, Global Change Biology 17, 1658 (2011).

[39] J. Laganiere, D. A. Angers, D. Pare, Global Change Biology 16, 439 (2010).

[40] K. M. Carlson, et al., Nature Climate Change 7, 63 (2017).

[41] G. Edwards-Jones, et al., Trends in Food Science & Technology 19, 265 (2008).

[42] D. Coley, M. Howard, M. Winter, British Food Journal 113, 919 (2011).

[43] P. Williams, et al., Conservation Biology 10, 155 (1996).

[44] J. F. Lamoreux, et al., Nature 440, 212 (2006).

[45] F. Albuquerque, P. Beier, PLoS One 10, e0119905 (2015).

[46] C. M. Roberts, et al., Science 295, 1280 (2002).

[47] W. Jetz, D. S. Wilcove, A. P. Dobson, PLoS biology 5, e157 (2007).

[48] B. International, H. of the Birds of the World, Bird species distribution maps of the world v6.0 (2016).

[49] NatureServe, IUCN, The IUCN Red List of Threatened Species v2016-1 (2016).

[50] N. Ramankutty, J. A. Foley, Global Biogeochemical Cycles 13, 997 (1999).

[51] X. Cai, X. Zhang, D. Wang, Environmental Science & Technology 45, 334 (2011).

[52] Y. Yang, Y. Luo, A. C. Finzi, New Phytologist 190, 977 (2011).

[53] P. Meli, et al., PLoS One 12, e0171368 (2017).

[54] Z. Fu, et al., Environmental Research Letters 12, 104004 (2017).

[55] W. Silver, R. Ostertag, A. Lugo, Restoration Ecology 8, 394 (2000).

[56] J. J. Gilroy, et al., Nature Climate Change 4, 503 (2014).

[57] K. J. Anderson-Teixeira, M. M. Wang, J. C. McGarvey, D. S. LeBauer, Global Change Biology (2016).

[58] L. Poorter, et al., Nature 530, 211 (2016).

[59] C. M. Johnson, D. J. Zarin, A. H. Johnson, Ecology 81, 1395 (2000).

[60] R. Houghton, Tellus B: Chemical and Physical Meteorology 51, 298 (1999).

[61] R. R. Dunn, Conservation Biology 18, 302 (2004).

[62] H. P. Jones, O. J. Schmitz, PLoS One 4, e5653 (2009).

[63] M. Curran, S. Hellweg, J. Beck, Ecological Applications 24, 617 (2014).

[64] D. Moreno-Mateos, et al., Nature Communications 8, 14163 (2017).

[65] R. L. Chazdon, et al., Conservation Biology 23, 1406 (2009).

[66] R. Crouzeilles, et al., Nature Communications 7 (2016).

[67] M. E. Bowen, C. A. McAlpine, A. P. House, G. C. Smith, Biological Conservation 140, 273 (2007).

[68] D. H. Dent, S. J. Wright, Biological Conservation 142, 2833 (2009).

[69] O. Hernández-Ordoóñez, N. Urbina-Cardona, M. Martínez-Ramos, Biotropica 47, 377 (2015).

[70] C. Sayer, J. Bullock, P. Martin, Biological Conservation 211, 1 (2017).

[71] J. M. R. Benayas, A. C. Newton, A. Diaz, J. M. Bullock, Science 325, 1121 (2009).

[72] K. D. Holl, T. M. Aide, Forest Ecology and Management 261, 1558 (2011).

[73] J. M. R. Benayas, J. M. Bullock, Ecosystems 15, 883 (2012).

[74] P. J. Seddon, C. J. Griffiths, P. S. Soorae, D. P. Armstrong, Science 345, 406 (2014).

[75] R. Crouzeilles, M. Curran, Journal of Applied Ecology 53, 440 (2016).

[76] R. Crouzeilles, et al., Science Advances 3, e1701345 (2017).

